# Assessing Pain in a Noninvasive Manner by Measuring Changes in the Microcirculation

**DOI:** 10.1101/2020.12.28.424536

**Authors:** Dana Shainshein, Louis Shenkman, Ahmed Khashan, Meir Bennun, Ilya Fine

**Affiliations:** Elfi-Tech Ltd., 2 Prof. Bergman St., Science Park, 76705 Rehovot, Israel; Tel-Aviv University, Israel; Meir Hospital, Kfar-Saba, Israel

**Keywords:** chronic pain, mDLS, dynamic light scattering, blood hemodynamic

## Abstract

The quantitative determination of the level of pain is one of the most challenging clinical problems. This article proposes a method for quantitative assessment of both acute and chronic levels of pain, based on the analysis of hemodynamic patterns measured using a non-invasive sensor. Hemodynamic characteristics were taken from the finger using a sensor measuring the dynamic scattering of light from the skin surface. Changes in hemodynamic parameters in patients with chronic pain were studied. One group of patients with chronic back pain required epidural injection for pain relief. The second group of patients had a Spinal Cord Stimulator implant which was switched off one hour before arriving at the clinic. Optical signals were collected before and after pain relief, either by epidural injection or by turning on the stimulator. Both groups reported their pain level using a standard numerical rating scale. Processing of the results showed that the changes in measured hemodynamic parameters corresponded in most cases to the changes in pain reported by patients following medical intervention. The results suggest that this new non-invasive measurement of pain can be used both for physiological studies and for monitoring various categories of patients suffering from pain.

## 1. INTRODUCTION

Assessment of pain is an important clinical problem for physicians caring for patients, especially in view of the current epidemic of opiate abuse. Since pain is a subjective experience, it is difficult for the physician to assess objectively the quality and intensity of the described pain. The ability to quantify pain would be an important contribution to the field of pain management. In this paper we discuss our efforts to develop an objective measurement system for quantifying the presence and intensity of pain, using an optical system based on dynamic light scattering to measure changes in the microcirculation.

The International Association for the Study of Pain defines pain as “an unpleasant sensory and emotional experience associated with actual or potential tissue damage” [1]. The sensory nervous system’s response to a harmful stimulus is the generation of pain. Pain is always a subjective, personal, and unpleasant experience. If the pain stimulus persists for a longer time pain becomes chronic. Chemical, mechanical, electric or thermal stimulations of nociceptors elicit a response from the central nervous system (CNS). Nociceptors require a minimum intensity of stimulation before they trigger a signal. Nociceptor stimulation results in various physiological responses and a subjective experience of pain [1].

Detection of nociceptive stimuli, such as surgical stimuli that actually or potentially cause tissue damage, relies commonly on the measurement of blood pressure or heart rate (HR) [2]. These measurements, however, may not detect all events or detect events with some delay [2]. New technologies have been developed to detect nociceptive events that rely on signals of the autonomic nervous system such as heart rate variability, pulse pressure, pupil diameter or skin conductance, or a combination of these signals [5, 9]. Most studies indicate that nociceptive indices based on these autonomic signals outperform BP and HR in the detection of nociceptive events [2–6]. Many patients with chronic pain do not obtain adequate relief or experience unacceptable side effects from existing treatments. Efforts to develop treatments that provide improved outcomes are therefore a priority for pain research and pain measurement.

Because of the subjective nature of the pain experience and the variation in pain threshold between individuals, assessing the severity of pain in the clinical setting is difficult. Currently, pain is assessed by observing the patient’s response and by a variety of subjective pain scores in clinical use. The development of an objective marker or methodology to assess pain would be a major advance in pain management [8]. In order to assess the degree of pain using non-invasive methods, it is necessary to identify measurable physiological parameters associated with pain. Ben-Israel et al [9] reported on the assessment of pain in a patient undergoing general anesthesia and found that the application of a multi-parametric approach that combined the heart rate variability indexes, skin conductance, and PPG perfusion could serve as a pain marker under anesthesia. However, as yet there have been no published works devoted to the measurement of chronic pain patients.

Another physiological manifestation of pain is a change in peripheral blood flow. The hemodynamics of the peripheral blood circulation is affected by the activity of the autonomic nervous system, which in turn responds to pain stimuli [20, 21]. Broens et al [10] described the use of a hemodynamic optical sensor (mDLS) to measure the response of peripheral blood flow to acute pain. The degree of response was shown to correlate with the level of acute pain. They were able to measure consistent, dose-related changes in the microcirculation in response to graduated pain stimuli, and to acute stressors. In the current study, we examined the effects of pain in two groups of patients suffering from chronic pain and whether pain relief can be measured with a hemodynamic optical sensor.

## 2. MATERIALS AND METHODS

### 2.1 The measurement system

The sensor technology used in this study relies on dynamic light scattering (DLS) [11]. A laser beam from a miniaturized probe (mDLS sensor, Elfi-Tech Ltd.) propagates through the skin and blood vessels, and the light scattered from the moving red blood cells (RBCs) creates a speckle pattern on the photodetectors. The sensor is made up of a 850 nm vertical cavity surface emitting laser (VCSEL) and two photodetectors (Fig. 1a-b). The sensor is connected to an electronic control unit (Elfor-S, Elfi-Tech Ltd.) that enables simultaneous measurement from two sensors. The control unit collects the data at 48 kHz and, using a computer interface, the raw data is stored in the memory and the processed signal is displayed in real time. The stored raw data is used for further signal processing.

**Fig. 1.**
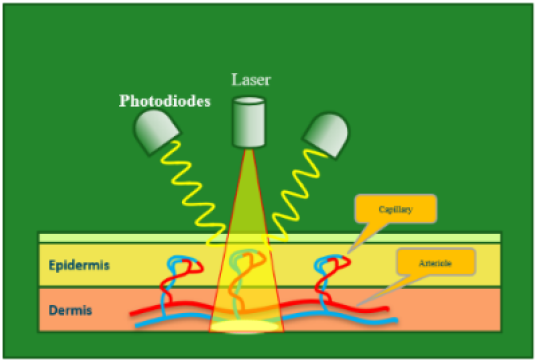
Schematic view of the sensor-skin interaction

**Fig. 1.**
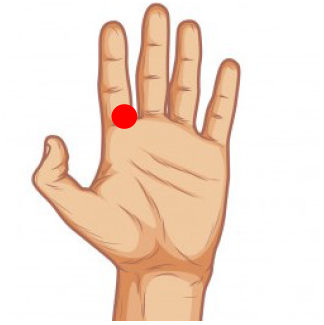
Illustration of the sensor’s location on a subject’s hand marked in red.

The measured signal originates from the light reflected from the moving red blood cells in the skin blood vessels. Different parameters related to the microcirculation are derived from this signal. Pain sensation, by acting through the autonomic nervous system, affects the microcirculation. [5–7, 19]. By detecting and analyzing the changes in the microcirculation we aimed to detect the presence of pain.

### 2.2 Patient Selection and measurements procedure

Two groups of patients aged 21 years or older of both sexes with chronic pain treated at the Meir Hospital Pain Clinic were enrolled in the study. The first group included 23 patients with chronic low back pain requiring the epidural injection of steroids and local anesthetic for pain relief. The second group was comprised of 5 patients treated with spinal cord stimulation for pain relief. All the patients were complaining of pain which had not responded to conservative treatment for at least 6 months. In both groups, patients were treated in the usual manner as required by their clinical status. Indications for treatment were based solely on standard clinical criteria. Their pain levels were assessed by the non-invasive sensor and by the administration of a numerical rating scale (NRS), the Short Form McGill Pain Questioner (SFMPQ), and the Patient Global Impression Scale (PGIS) rating for improvement with treatment during several time points of their treatment. The study was approved by the local Human Ethics Committee (Meir Hospital, Kfar Saba, Israel) and was registered at www.clinicaltrials.gov under number NCT01912118. All patients gave oral and written informed consent before enrolment into the study.

### 2.3 Study protocol

#### a) Epidural injection group

Patients in this group had chronic low back pain and were referred to the Pain Clinic for treatment with spinal injections (epidural, transforaminal, facet joint blocks) under fluoroscopy. These patients arrived at the pain clinic and initially were screened by the nurse. During the initial screening, patients were asked to note their pain level on an NRS scale (from 0 to 10) and to complete the SFMPQ and PGS assessments. The sensors were then placed on the finger root and a recording was taken for 5 minutes. The patients then were taken to the radiology suite where they received a spinal injection by the pain specialist. After the injection, the patient returned to the clinic for an observation period of 30 to 60 minutes. After resting for 30 minutes, an additional 5-minute device recording and NRS scale, SFMPQ, PGIS assessments were carried out. 2 weeks after the injection the sensor was again attached to the finger for the third assessment. Additional NRS scale, SFMPQ and PGIS assessments, and 5 minutes measurement with the sensors were performed.

#### b) Spinal cord stimulation patients

Patients attending the Pain Clinic who were being treated with Spinal Cord Stimulation (SCS) were studied. Patients had the implant inserted two months or more prior to the study. The patients were requested to switch off the stimulator one hour before arriving to the clinic. During the clinic visit, patients were asked to note their pain level on a NRS scale and SFMPQ. The sensor was then placed on the finger and recordings were acquired for 5 minutes. After the initial assessment the stimulator was switched on for 20-60 min at its previous setting. A second measurement with the sensor was obtained and the NRS, SFMPQ and PGIS assessments were performed again.

### 2.4 Measurement procedure

The signals were collected while a subject was sitting in a semi-reclining position and immobile. mDLS sensor was positioned on the skin and gently fixated with adhesive tape. The sensor was placed on the palmar aspect of the root of the left index finger (Fig. 2).

## 3. TECHNOLOGICAL BACKGROUND

### 3.1.1 Hemodynamic Indexes

The relative movement of RBC’s particles in the blood vessels is described by a velocity profile *v(r,t)* of blood flow. In a very simplified case, for the vessel of radius *R,* axis-symmetric velocity profiles, *v(r,t)* can be described in cylindrical coordinates by this empirical relationship:

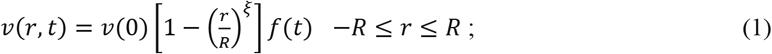

Where *v*(0) - is maximum velocity at the center position r=0 and R is the radius of the vessel, *f(t)* is a periodic function of heart beat frequency, which is driven by the difference between systolic and diastolic pressure wave and it is time phase-shifted with respect to the cardiac cycle and represents the degree of blunting. The velocity gradient or shear rate is determined by the diameter of vessels and the average velocity of the flow. If we have a blood vessel with velocity distribution *v(r, t)* where r is the cross-session of the vessel, the shear rate *γ* is defined by:

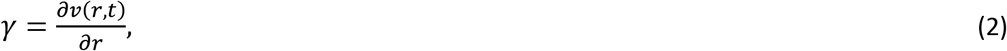

The average shear rate in the vessel can be estimated by:

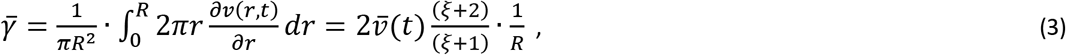

From (3) we can see that the mean velocity and the radius of the vessels determine the average blood shear rate. The high shear rate corresponds to the fast flow and the small vessel diameter and low shear rate corresponds to the slow flow and a large diameter of the vessels. Thus, the shear rates distribution can be associated with different types of blood vessels or different regions inside the vessels. The lowest shear rate values correspond to the RBCs located mostly near the walls or flowing through the narrow capillary blood vessels and their decay function is dominated by the Brownian movement statistics. The very high shear flow is related to the large capillary vessels and arterioles where the pulsatile component of the signal becomes to be prominent. Most of the parameters that determine the shear rate are not constant and the shear rate is modulated not only by the heart-beat frequency but in a more complex manner.

The use of DLS or laser Doppler makes it possible to extract the information related to the shear rate modulation and associated blood flow parameters by analyzing the signal that is obtained from the skin surface. The DLS parameter that is directly related to the shear rate value is the decay time of the autocorrelation function of the signal. For the measured signal I(t) we can derive the autocorrelation function g by:

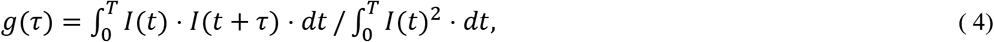

In the case where Brownian motion is negligible and the light intensity fluctuations are contributed mainly by the shear rate, we can use the following expressions [11]:

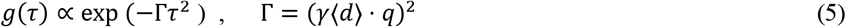

where *q = 2ksin(θ/2)* where - *θ* is scattering angle, k is wavelength number, <d> is the effective distance across the scattering volume in the direction of the velocity gradient. When we consider the blood vessels located inside the tissue, the value of q should be calculated after averaging over all directions. For blood flow analysis, it is more convenient to present the signal in terms of spectral density for any given value of shear rate. The relationship between the autocorrelation function and power spectrum for (5) can be given by[13]:

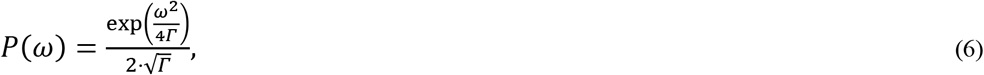

Practically, when the laser beam is reflected from the surface of the skin, a variety of blood vessels of different diameters and different shear rates contribute to the formation of the signal. Therefore, it should be taken into consideration that the measured signal is a superposition of the signals that originated from different vessels.

In [12] the so-called hemodynamic index (HI) was introduced. This HI represents the integral of the power spectrum taken in a predefined spectral band. Thus, each HI is determined by the frequencies band ω_1_ and ω_2_:

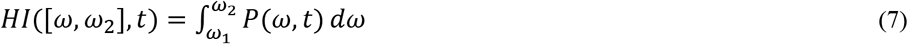

According to the physiological interpretation, HI corresponds to a certain shear rate values and represents a subtype of a blood vessel. It is possible to distinguish between large vessels such as arteries and arterioles and small vessels such as capillaries or venules, for instance, by observing a pulsatile pattern resembling the blood pressure wave in HIs that are associated with pulsatile blood vessels (fig. 3).

**Fig. 3.**
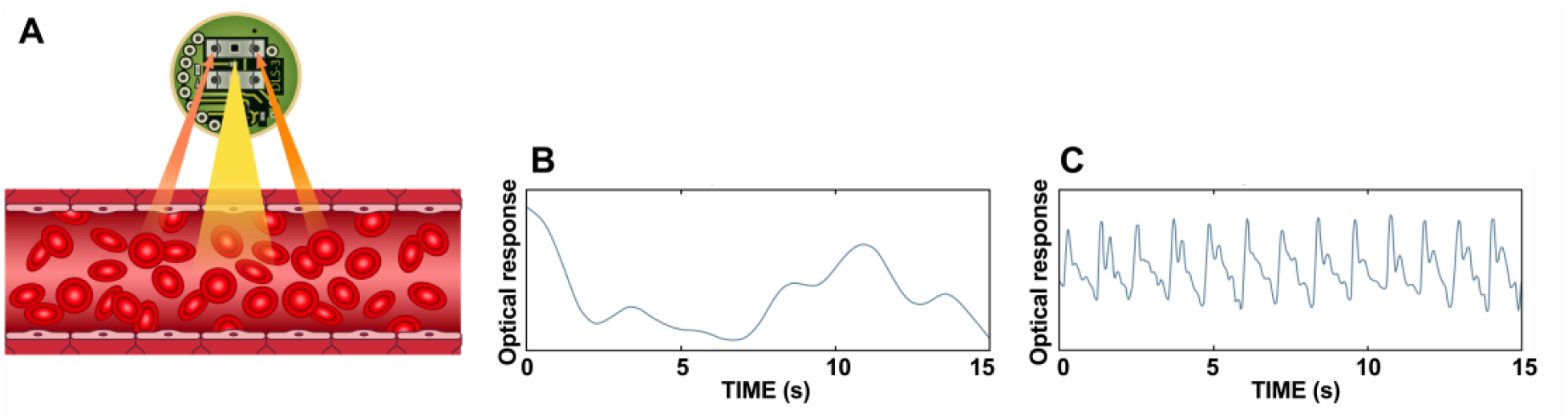
Schematic overview of the miniature dynamic light scattering (mDLS) technique used in this study. The small (diameter 1 cm) mDLS sensor radiates laser light through the skin. The light reflected from flowing red blood cells is collected via two optical sensors (A) and further analyzed. Non-pulsatile (B) and pulsatile (C) signal are derived from the fluctuations in the intensity of the reflected optical signal through power spectrum analysis

While HI is an absolute parameter that may vary between trials due to slight changes in sensor location or proximity to the skin, the relative HI (relHI) is a normalized parameter defined as:

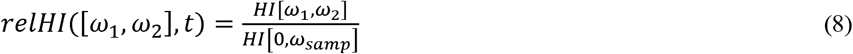

where *ω_samp_* is the sampling frequency of the measured signal, therefore *HI*[0, *ω_samp_*] represents the total energy and can also be defined as the perfusion, as it contains all the frequency bands. The variations in relHI following physiological events can be compared between different measurements.

An additional parameter that is extracted from the mDLS signal is the relative blood flow velocity. This parameter is equivalent to the statistical representation of the skin blood velocity which can be derived from Doppler flowmetry [17]. Formally, the equivalent skin blood velocity parameter can be defined as the normalized first moment of the power spectrum of the mDLS sensor signal. We term this parameter the relative blood velocity (RBV).

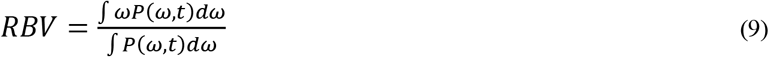

### 3.1.2 Oscillatory Indexes

The blood flow is subject to oscillations at different frequencies (pf), each of which is caused by different physiological processes [14,15,16]. The highest band of the physiological frequencies (8-2 Hz) is associated with the pulse component. The next range of frequencies (0.12-0.4) is associated with the respiratory component. Frequency Interval from 0.06 to 0.15 Hz is related to the blood pressure regulation and is known as the myogenic response. The frequency interval below 0.06 Hz is related to neurogenic and endothelial activity. Thus, each HI can be characterized by related patterned of oscillations. These patterns are called Oscillatory Hemodynamic Indexes (OHI) [13]. OHI is obtained from Power Spectrum (PS) of HI(ω_1_,ω_2_, *t*). PS of HI is analyzed from pf=0 to pf=2Hz.

OHI for each band should be defined. For example, for the pulsatile component pf ranges from 0.8 to 2 Hz or for the myogenic response pf is between 0.5 and 1.5 Hz. OHI is normalized to the full range of frequencies pf (0 −2 Hz):

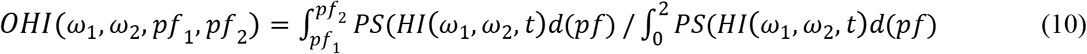

Figure (4) shows examples of spectral behavior (OHI) for four hemodynamic indices HI. Besides, we can see examples of HI spectrum of chronic patient subjects for two HI components - non-pulsatile (0-1Khz) and pulsatile

**Fig. 4.**
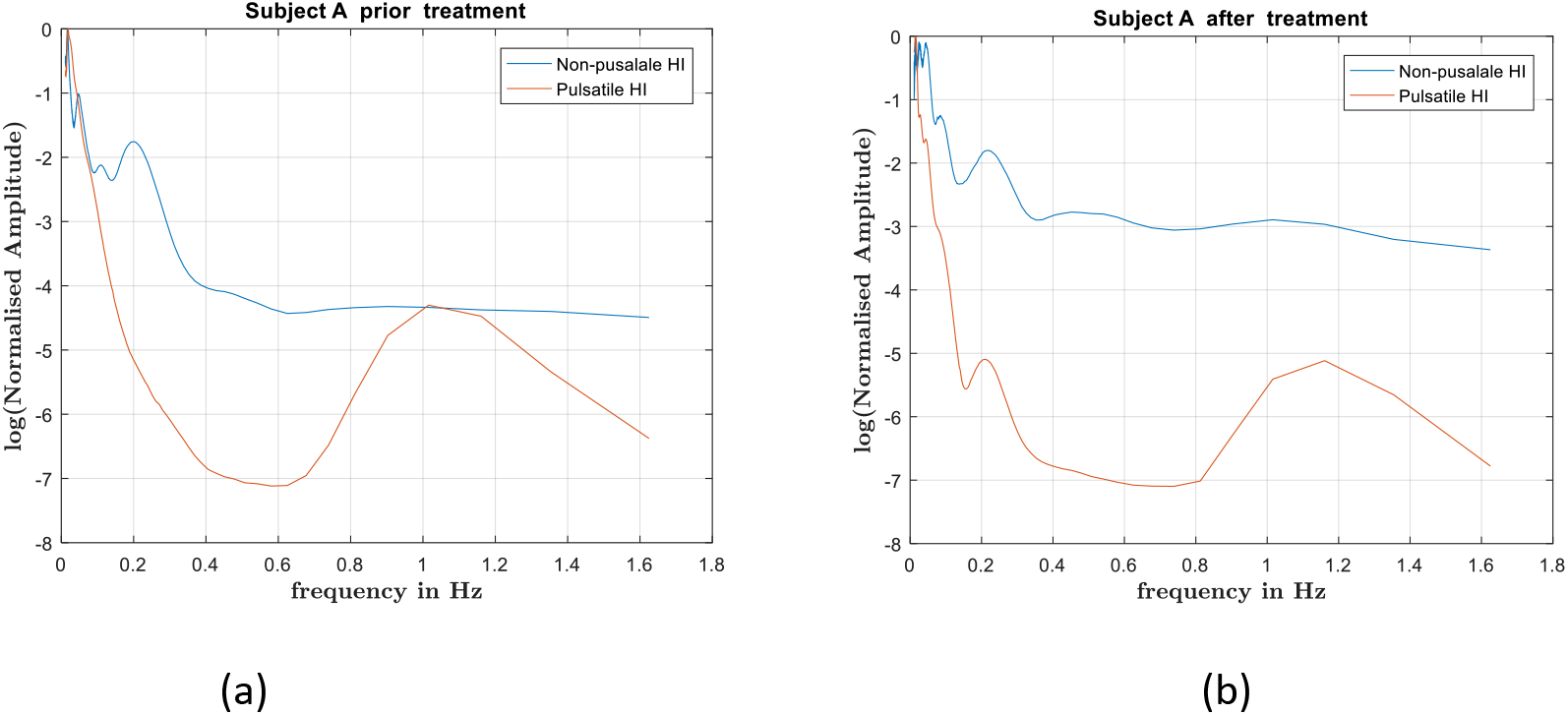
Spectral behaviors for non-pulsatile and pulsatile HI for a chronic pain subject before epidural (a) and two weeks after epidural (b)

The OHI parameters are closely related to characteristic oscillation arising from central regulatory mechanisms [14,15,16] and result from blood pressure waves and rhythmic oscillations in vessel diameter or vasomotions. Another mechanism of blood flow oscillation evolves the response of endothelial cells[22]. For example, arterial smooth muscle relaxation is partially mediated by endothelial-dependent mechanisms, which involve the release of nitric oxide. Thus, physiological changes occurring in the body are measured in terms of two variables - hemodynamic indices (HI) which are related to the type of blood vessels and oscillatory indices (OHI) which are associated with different dynamic physiological mechanisms.

Rationale: Our goal was to determine the significance of the various hemodynamic measures in relation to chronic pain. To this end, we tested the responses of four hemodynamic measures derived from the optical signal of the mDLS sensor: relHI, OHI, RBV and perfusion at two separate levels of pain and to determine if differences can be detected. These hemodynamic parameters are directly related to autonomic nervous system activity[14], and could potentially be used for detecting (and quantifying) the autonomic behavior during chronic pain.

## 4. RESULTS AND DISCUSSION

The hemodynamic characteristics reported here represent the differences between the average value of the calculated parameters, taken prior to medical intervention, and after. In the Epidural injection group, the differences were before and after epidural injection, and in the Spinal Stimulation group, they were before and after turning on the stimulator. The differences between reduced and increased pain values were calculated for each of the parameters derived from the measured signal. Delta (relHI) was calculated for the HI representing the blood flow in the small blood vessels, such as capillaries or venules (frequency band of 0-1 kHz). Delta (OHI) was derived from HI representing oscillating high shear rates of greater vessels, such as arterioles (frequency band 1-24 kHz).

We analyzed two types of physiological responses: the short-term response and the long-term response. For shortterm response, reduced pain refers to (1) measurement 30 minutes after epidural injection or (2) measurement with SCS device turned on. Increased pain values refer to (1) measurement before epidural injection or (2) measurement 30 minutes after the SCS device was turned off. For long-term response, reduced pain refers to measurement 2 weeks after epidural injection. The data was analyzed using a paired-t-test with p-values < 0.05 considered significant.The short-term response was compared to the results of a previous study [10] where the hemodynamic response to acute pain was measured. In the study of acute pain hemodynamic responses were taken as a response to the induced pain. In this study we observed the hemodynamic responses to pain relief.

In the SCS group one subject was measured twice (measurements 2,3 in the Figure 5). In the epidural group some data could not be used for analysis due to low quality; therefore, in the short-term response epidural group there are only 15 patient and in the long-term response epidural group there are 17 patients.

**Fig. 5.**
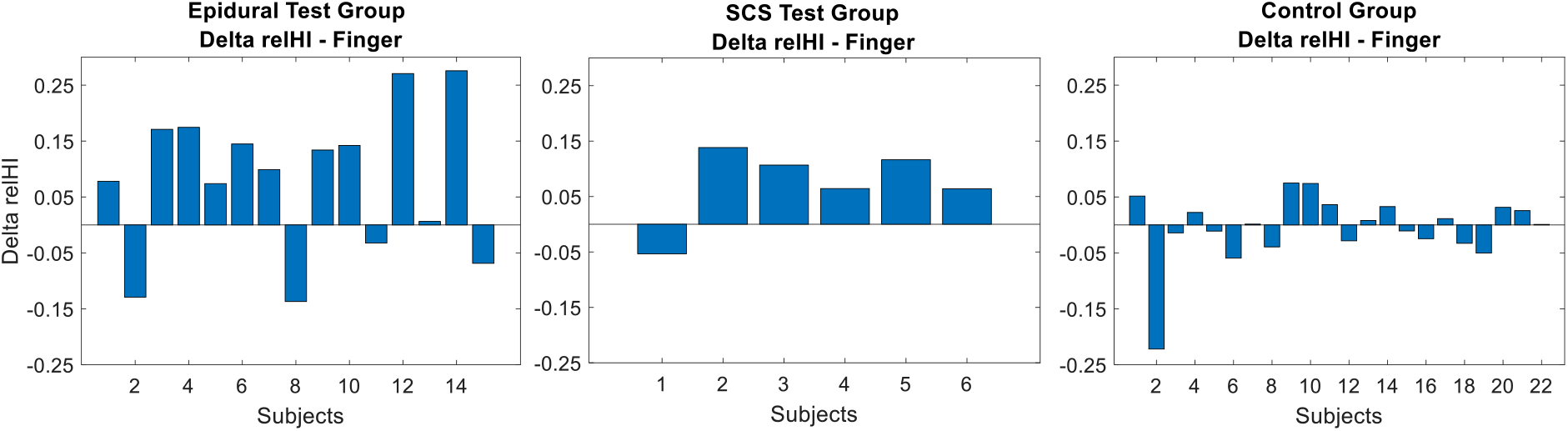
A statistical representation of Delta (RelHI) from capillary blood vessels response to short-term increased pain. Values are the difference between mean relHI during reduced pain subtracted from increased pain.

### 4.1 SHORT-TERM RESPONSE

#### 4.1.1 Relative hemodynamic index response

An increase in relHI originated from the capillary blood was observed for most patients, in both epidural and SCS groups in the short-term response (Figure 5, Table 1). This finding is consistent with the similar acute paint response measured in [10].

**Table 1.**
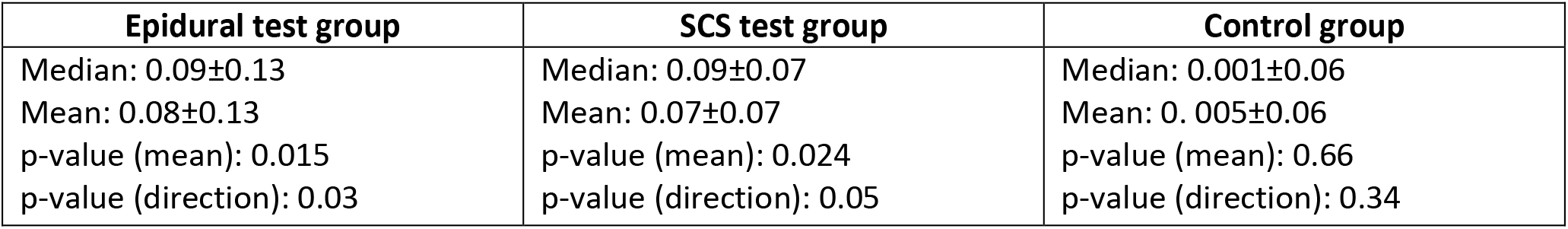
Summary of the relHI response in each group to short-term increased pain. Data presented is median/mean ± SD measured from all the patient’s delta mean values.

#### 4.1.2 Relative blood velocity response

The RBV response was significant only in the SCS groups, with a decrease in RBV during increased pain (Figure 6, Table 2).

**Fig. 6.**
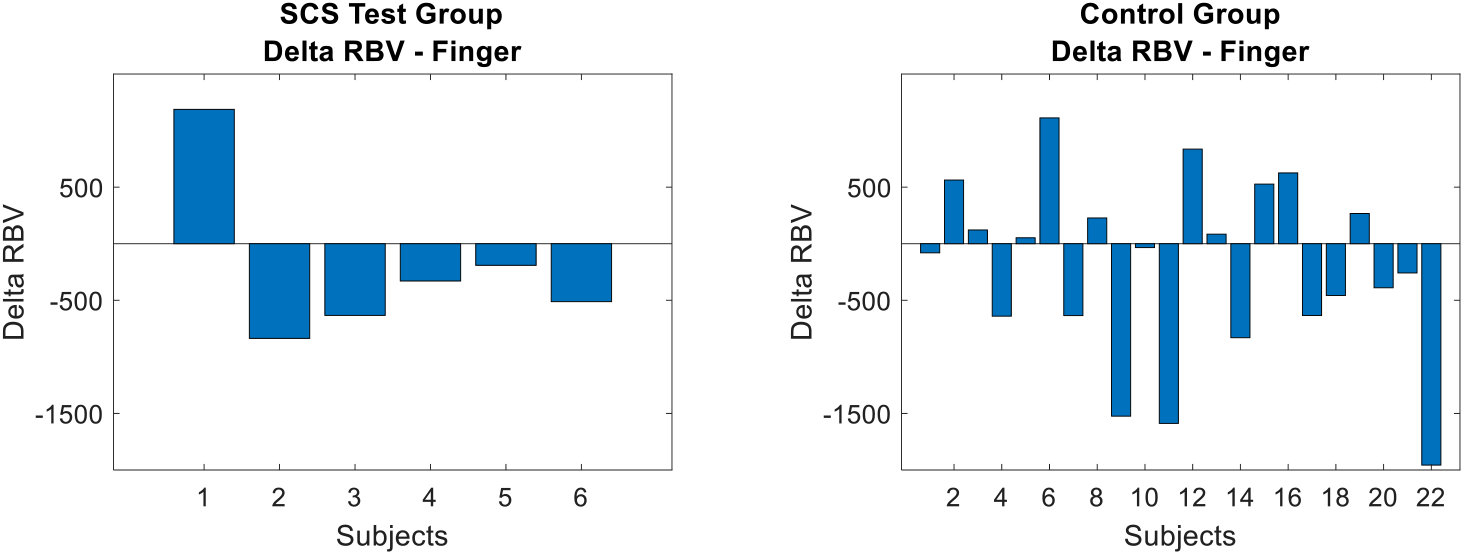
RBV response to short-term increased pain. Values are difference between mean RBV during reduced pain subtracted from increased pain.

**Table 2.** Summary of the RBV response to short-term increased pain measured from the finger in the SCS group. Data presented is median/mean ± SD measured from all the patient’s delta mean values.

| SCS Test group – RBV from finger | Control group – RBV from finger |
| --- | --- |
| Median: -420±725<br>Mean: -219±725<br>p-value (mean): 0.25<br>p-value (direction): 0.05 | Median: -57±790<br>Mean: -209±790<br>p-value (mean): 0.11<br>p-value (direction): 0.34 |

#### 4.1.3 Total perfusion response

A response to short-term pain was also observed in the perfusion parameter only in the SCS group (Fig.7 and Table 3).

**Fig. 7.**
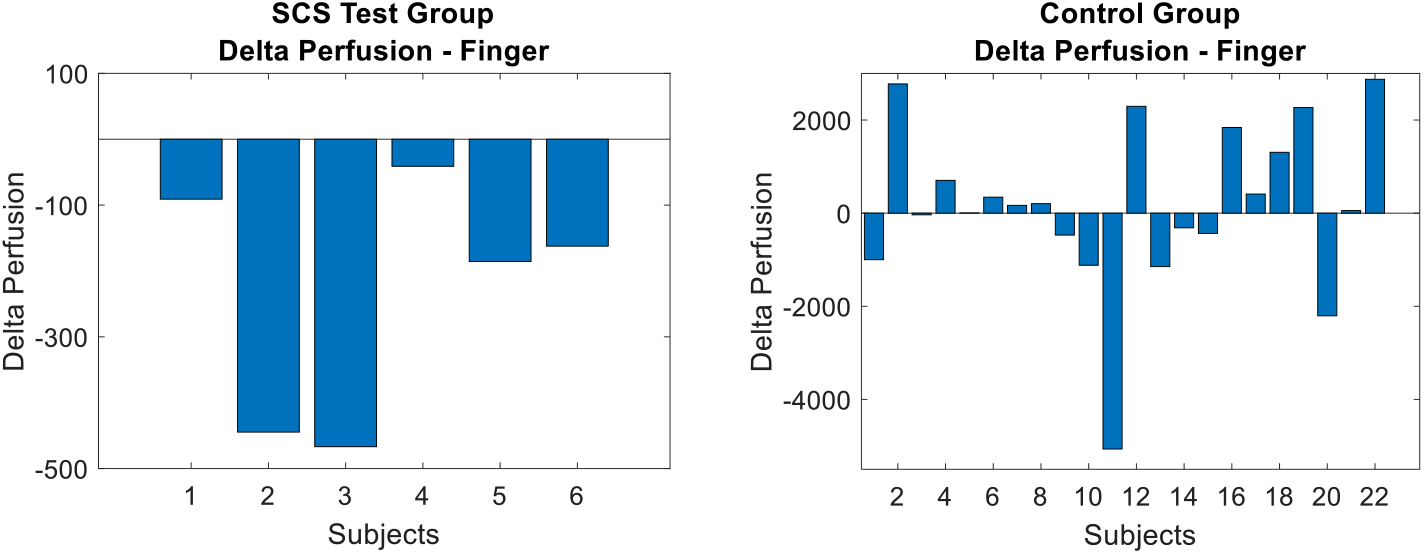
Perfusion response to short-term increased pain is measured in the SCS group. Values are the difference between mean perfusion during reduced pain subtracted from increased pain.

**Table 3.** Summary of the perfusion response to short-term increased pain. Data presented is median/mean ± SD measured from all the patient’s delta mean values.

| SCS Test group – Perfusion from finger | Control group – Perfusion from finger |
| --- | --- |
| Median: -420±725<br>Mean: -219±725<br>p-value (mean): 0.01<br>p-value (direction): 0 | Median: -57±790<br>Mean: -209±790<br>p-value (mean): 0.11<br>p-value (direction): 0.34 |

### 4.2 Long-term response

#### 4.2.1 Oscillatory hemodynamic index response

A significant increase in pulsatile OHI was observed for most chronic pain patients, 3 weeks after the treatment by the epidural (see Fig. 8 and Table 4).

**Fig. 8.**
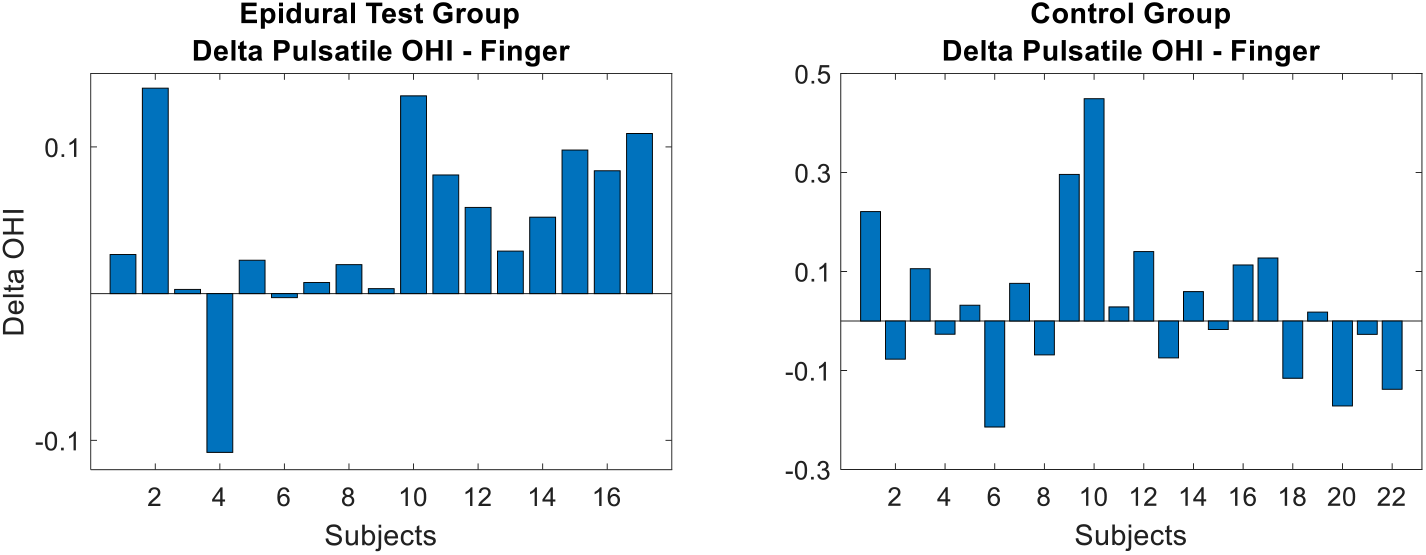
Pulsatile OHI response mesaured from finger root to long-term increased pain. Values are difference between mean OHI during reduced pain subtracted from increased pain. In the test group the majority of the subjects displayed an increase in pulsatile OHI 2 weeks after the procedure.

**Table 4.** Summary of the pulsatile OHI response to long-term increased pain. Data presented is median/mean ± SD measured from all the patient’s delta mean values.

| Epidural test group | Control group |
| --- | --- |
| Median: 0.03±0.06 | Median: 0.02±0.16 |
| Mean: 0.045±0.06 | Mean: 0.03±0.16 |
| p-value (mean): 0.004 | p-value (mean): 0.84 |
| p-value (direction): 0.0001 | p-value (direction): 0.66 |

#### 4.2.2 Total perfusion response

A significant response to long-term pain response was also observed in the perfusion parameter, 2 weeks after the procedure (Fig.8 and Fig.9).

**Fig. 9.**
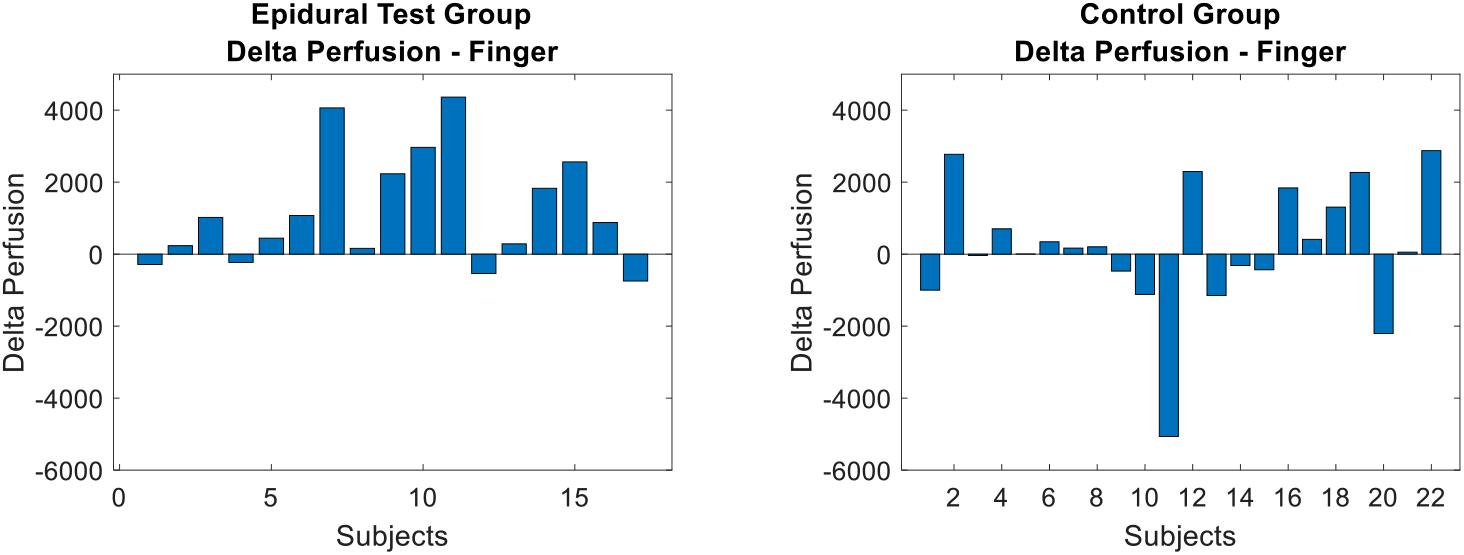
Perfusion response mesaured from finger root to long-term increased pain. Values are difference between mean perfusion during reduced pain subtracted from increased pain. In the test group the majority of the subjects displayed an increase in perfusion 2 weeks after the procedure.

| Epidural test group – Perfusion | Control group – Perfusion |
| --- | --- |
| Median: 877±1567 | Median: 111±1778 |
| Mean: 1195±1567 | Mean: 157±1778 |
| p-value (mean): 0.003 | p-value (mean): 0.34 |
| p-value (direction): 0.01 | p-value (direction): 0.20 |

The perfusion changes also correlate with the subjective NRS, SFMPQ and PGIS assessments as it is shown in Figure (10). The three pain assessment methods were summed for the analysis and are referred to as “Pain Score”.

**Fig. 10.**
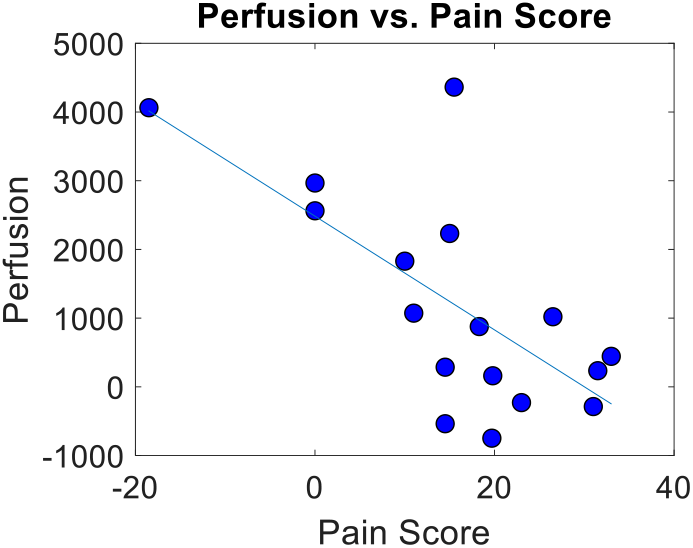
Perfusion from finger root plotted in relation to each subject’s reported change in pain level (pain score). A linear regression line was drawn, with RMSE = 0.001 and adjusted R^2^ = 0.44.

A linear relationship was observed between perfusion measured from finger root and the subject’s reported change in pain score (Figure 10).

### 4.3 Discussion

Because the sensation of pain is a subjective phenomenon, it is very difficult to assess its intensity in an objective manner. While severe pain may be accompanied by physical manifestations such as facial grimacing, guarding of the painful area, and changes in pulse rate or blood pressure, it is often not possible to determine the degree of pain which the person is experiencing by observation. Pain assessment in non-verbal individuals and in infants is also an important clinical problem. More pressing today in the face of the current opiate overuse epidemic are individuals who claim they are in severe pain and request analgesics. An accurate, reproducible, and objective measure of pain severity would be a boon for pain management protocols.

The autonomic nervous system is engaged during the elicitation of pain, and changes in autonomic nervous system function are reflected in the microcirculation. Using a hemodynamic optical sensor, we were able to assess the presence and severity of acute pain in an experimental setup [10].

**Fig. 11.**
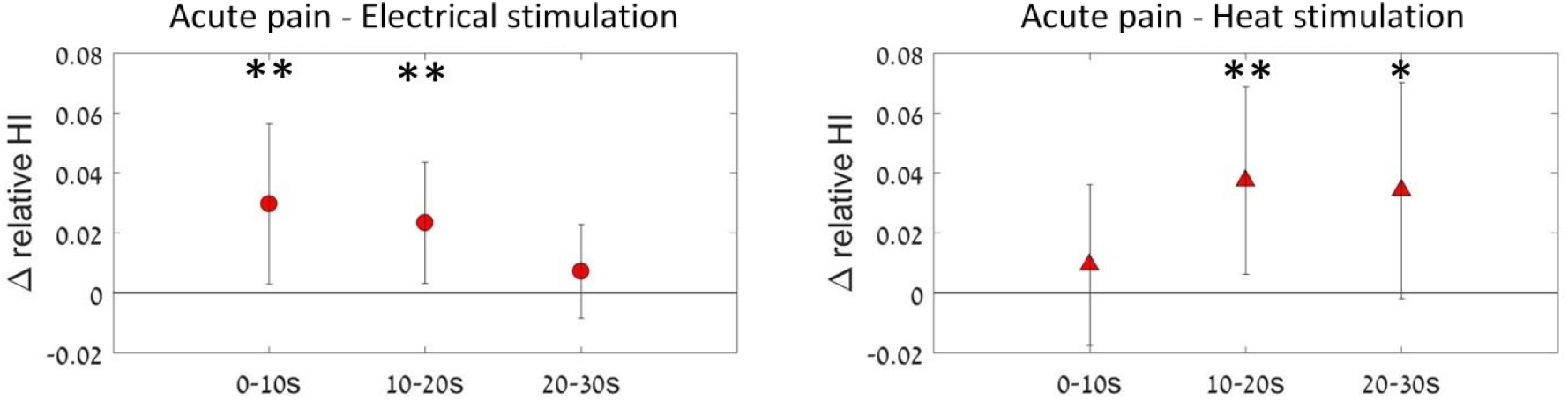
RelHI response originated from capillary blood vessels taken from acute pain study6. Temporal profile of the change in relHI during electrical and heat stimulation (acute pain) at different stimulus intensities as measured from left index finger. The data are divided into three time windows, 0-10 s, 10-20 s and 20-s, of stimulus time. Values are mean ± SD. * p < 0.005, ** p <0.0005.

In the current study, we applied the same experimental methodology to assess the presence and severity of chronic pain in two separate patient groups. In the first group, patients with chronic low back pain requiring epidural anesthesia were assessed before and after the procedure, when pain intensity was expected to be lower. In the second group, patients utilizing a spinal cord stimulator for pain relief were studied after the stimulator was turned off and the pain was expected to be of higher intensity, and then after restarting the stimulator with expected relief of pain. We determined the relative blood velocity response and total perfusion response and two levels of pain, and then we assessed the long-term response of these hemodynamic indices.

Our results show that changes in pain levels are reflected in hemodynamic changes and in the microcirculation. The relHI at the finger root increased in most patients in both the epidural and SCS groups, similar to the changes we observed in our study of acute pain (10).

Regarding the relative blood velocity response, the epidural group and the SCS group showed a decrease in RBV during increased pain (Figure 6, tables 2). A similar response was observed in the previous acute pain study. When tested several weeks later, both groups showed a significant increase in the perfusion parameter at two weeks, with a linear relationship between the subject’s reported change in pain score. A significant response to long-term pain was also observed in the perfusion parameter 2 weeks after the procedure. An inverse linear relationship was observed between perfusion measured from the finger root and the subject’s reported change in pain score (Figure 10). The blood flow short term response to pain is consistent with the well-known fact that pain activates the sympathetic nervous system, increase muscle sympathetic nerve activity, increase in the sympathetic drive that can lead to peripheral vasoconstriction and arterial blood flow reduction (19).

The most statistically significant pain markers have been obtained in a comparative analysis of the oscillatory blood flow indices (OHIs). The periodic oscillations in cutaneous blood flow can be modulated by the vascular endothelium, and may be assessed by the difference in effects of endothelium-dependent and endothelium-independent vasodilators like nitric oxide on the blood flow oscillations of this frequency. Indeed, there is good evidence that cytokines and nitric oxide (NO) play an important role in central and peripheral modulation of nociception [18].

## CONCLUSIONS

These results indicate that chronic pain influences the microcirculation, and open the possibility of objectively measuring pain. Of all the hemodynamic indices measure, we found that the Oscillatory Hemodynamic Index (OHI) was the most robust. It is possible that by combining several indices, a stronger correlation between pain level microvascular hemodynamic changes may be established.

